# eccDNA purification, full-length sequencing, and genomic mapping

**DOI:** 10.1101/2022.06.15.496131

**Authors:** Yuangao Wang, Meng Wang, Yi Zhang

**Affiliations:** Howard Hughes Medical Institute, Boston, MA 02115; Program in Cellular and Molecular Medicine, Boston Children’s Hospital, Boston, MA 02115; Department of Genetics, Boston, MA 02115; Harvard Stem Cell Institute WAB-149G, 200 Longwood Av., Boston, MA 02115

## Abstract

Extrachromosomal circular DNA (eccDNA) has been discovered for more than half a century^1^. However, its biogenesis and function have just begun to be elucidated. One hurdle that prevented our understanding of eccDNA is the difficulty in obtaining pure eccDNA from cells. The current eccDNA purification methods mainly rely on exonuclease digestion after alkaline lysis. Due to its low abundance and heterogeneity in size, the current eccDNA purification methods are not efficient in obtaining pure eccDNA. Here, we describe a new 3-step eccDNA purification (3SEP) procedure by adding a new step that allows circular, but not linear, DNA that escaped from exonuclease digestion, bound to silica beads in solution A. Our method allows eccDNA purification with high purity and reproducibility. Additionally, we developed a full-length sequencing and genomic mapping method by combining rolling cycle amplification (RCA) with Nanopore sequencing, through which the consensus sequence of multiple tandem copies of eccDNA contained within the debranched RCA product is derived and mapped to its genomic origin. Collectively, our protocol will facilitate eccDNA identification, characterization, and has the potential for diagnostic and clinic applications.

## Introduction

EccDNAs were known for their broad existence in almost all cell lines, tissues and organs across species since their first description in 1964^1^. Robust eccDNA purification methods are crucial for identifying their genomic origin and understanding their biogenesis. However, their size heterogeneity and low abundance relative to their linear chromosomal counterpart make obtaining pure eccDNAs extremely difficult. Hirt^2,3^, Alkaline lysis^4^, exonuclease digestion^5-7^, buoyant density^2,3,5,8^, and their combinations have been used for eccDNA purification. In general, crude DNA circles are extracted from biological samples by Hirt^2,3^ and alkaline lysis^4^, by which large size chromosomal DNAs are co-precipitated and removed with proteins. Then, linear contaminant DNAs within the crude circular DNA are digested with an exonuclease, such as ATP-dependent Plasmid Safe DNase (P.S. DNase)^6^, exonuclease V (RecBCD)^7^, or exonuclease III^9^. However, two precautions have to be taken when using exonuclease treatment. First, the specificity and efficacy of the exonuclease used have to be examined because trace amount of endonuclease activity in the exonuclease can cause eccDNA loss, particularly considering the extended and multiple rounds of exonuclease digestion widely used; Second, the exonuclease activity can be inhibited or blocked by protruding termini^10^, certain abnormal and damaged nucleotides^11^, cross linkages^12^, special structures^13^ etc., or their digestion products^6^. Thus, treatment of the crude circular DNA with exonuclease alone may not be sufficient for obtaining pure eccDNA, as demonstrated by a previous study using alkaline lysis and extended exonuclease digestion^14^. Although the crude circular DNA from Hirt, alkaline lysis or even exonuclease treatment could be further purified by cesium chloride-ethidium bromide (CsCl-EB) gradient ultracentrifuge^3^, given its laborious nature ^15^, it does not satisfy the needs of most studies.

Here we use a reagent that is not related to DNA denaturation or exonuclease and is able to selectively bind and recover circular DNA on silica beads while leaving linear DNA in solution, to further enrich eccDNA. Eventually, we developed a robust 3-step eccDNA purification (3SEP) technique (**Fig.1a**) ^16^. In the first step, the crude DNA circles are firstly extracted with buffered (pH11.8) alkaline lysis, a condition that causes less circular DNA loss than the conventional 0.2 M sodium hydroxide^17^. In the second step, linear DNA in the crude circular DNA is reduced by the treatment of ATP-dependent Plasmid Safe DNase, meanwhile the circular mitochondrial DNA is linearized by restriction digestion of an 8-bp cutter PacI to make our method compatible for eccDNA extraction from whole cells. Finally, a solution (Solution A) is used to recover eccDNAs selectively by their capacity of binding circular DNA to magnetic silica beads in this solution while leaving the contaminating linear DNA in the solution.

The 3SEP protocol is designed for eccDNA purification from cultured mammalian cells. To adapt the protocol for eccDNA purification from tissue samples or other type of samples, proper sample processing, such as cell dissociation or tissue grinding, may be needed before starting 3SEP.

To obtain the full-length sequence of eccDNAs, we use Oxford Nanopore to sequence the long DNA molecules that are generated by debranching the rolling circle amplification (RCA) products of the purified eccDNAs (**Fig. 1b**). Since the nanopore reads of the long DNA molecules are consisted of tandem multiple copies of the templated eccDNA, the consensus sequence derived from these tandem copies of a long read represents the full-length sequence of the templated eccDNA (**Fig. 1b**). To determine the genomic origin(s) of each eccDNA, each Nanopore read was mapped to the reference genome and divided into subreads based on the location of each mapped portion. If all the subreads are mapped to a single locus, this suggests that the eccDNA is from self-ligation of one genomic fragment. If the subreads are mapped to two or more different loci in the same or different chromosomes, the eccDNA is formed by ligation and circularization of multiple genomic fragments. To determine the boundary (joining ends) and consensus sequence for both single- and multiple-fragments eccDNA, we developed a unified threading method to reconstruct the structure of eccDNA based on the order and location of mapped subreads in each Nanopore read (Box 1). We implemented the eccDNA calling method in a single script and provided an integrated computational pipeline from raw sequencing signal processing to generating final eccDNA consensus.

**Figure 1.**
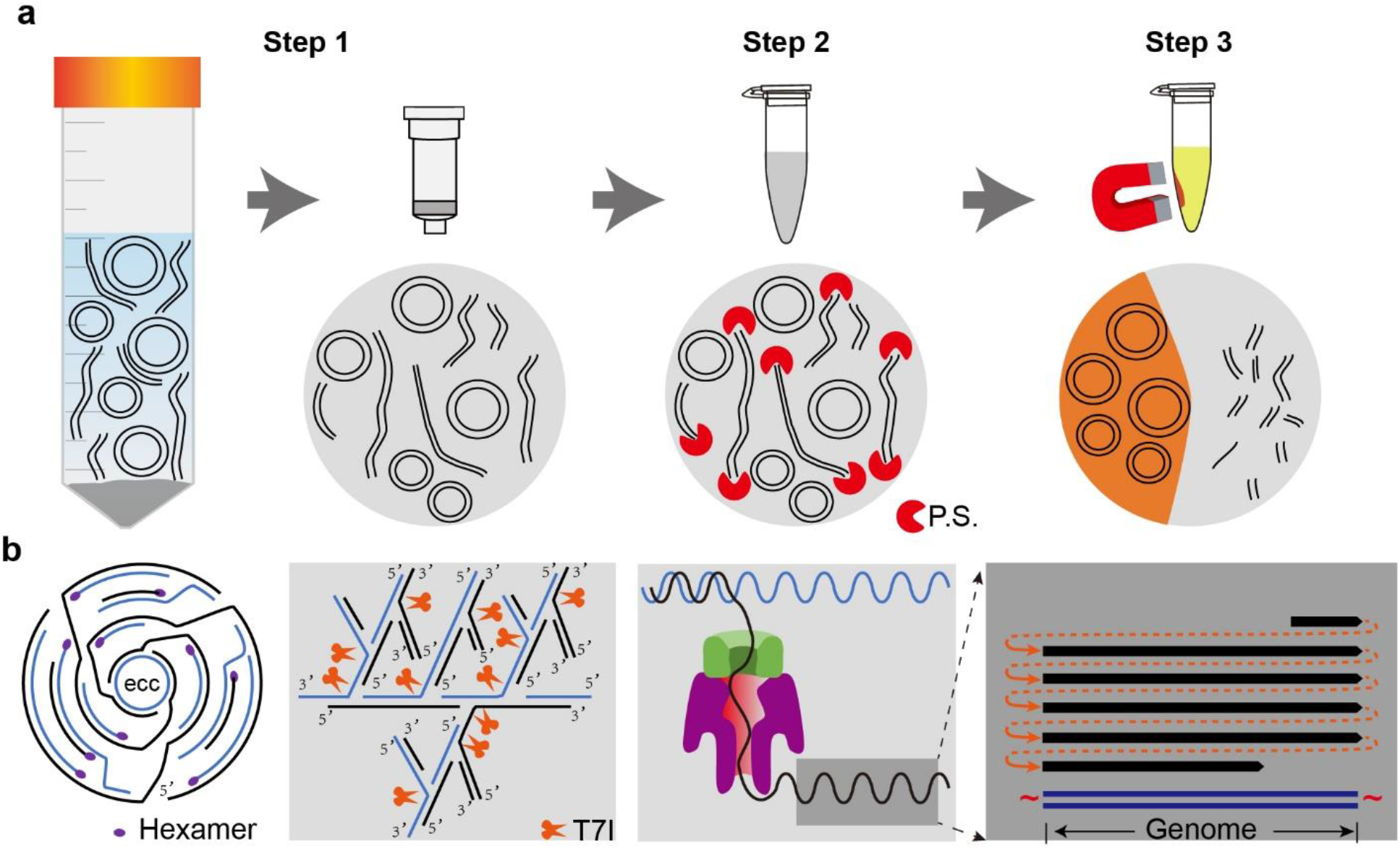
Scheme of the eccDNA purification and sequencing procedure. **a)** The 3SEP scheme. Step 1, crude DNA circles are extracted from whole cells with a buffered alkaline lysis at pH 11.8, bound and eluted from silica column; Step 2, linear DNA and PacI linearized mitochondrial DNA are digested with Plasmid-Safe (P.S.) DNase; Step 3, eccDNAs are selectively recovered with magnetic beads and solution A. **b)** EccDNA sequencing procedure. EccDNAs are firstly amplified by rolling cycle amplification and debranched with T7 endonuclease I (T7I), then read through by Oxford Nanopore. The full-length sequence of the eccDNA is called as consensus sequence of multiple tandem copies within long read.

Compared to the previous eccDNA purification methods^14^, our method not only results in eccDNA of high purity, as revealed by microscopy imaging (**Fig. 2b** and **d, right panel**), but also saves time, as the purification procedure can be completed within one day instead of a week^18,19^. Additionally, given its high reproducibility, our method can be used for evaluating the eccDNA yield by visualizing in an agarose gel (**Fig. 2a** and **c**) as well as in our previous study ^16^. Additionally, Solution A and magnetic silica beads described in Step 3 of 3SEP could be applied to enrich circular DNA of other types, such as mtDNA, viral DNA etc… Furthermore, by Nanopore sequencing, our method can also be used to reconstruct eccDNA full-length sequences and to reveal the genomic origins, including the self-ligation of a single genomic fragment and multiple-ligations of fragments from different genomic regions. Our computational method could calculate the number of full-passes, which measures the reoccurrence of the same structure (concatemer) for rolling-circle amplified circular DNA. A minimum of two full-passes ensures the circularity of the original DNA fragments as linear DNA or random jumping ones would not repeat the same structure for two or more times. To avoid artificial concatemers generated from branching and template-switching during phi29 amplification, we only kept eccDNAs with all fragments in the same locus mapped to the same strand, since artificial concatemers generated from branching and template-switching would map to different strands in the same locus. Thus, our purification and computation method together ensure the high reliability of the identified eccDNAs.

**Figure 2.**
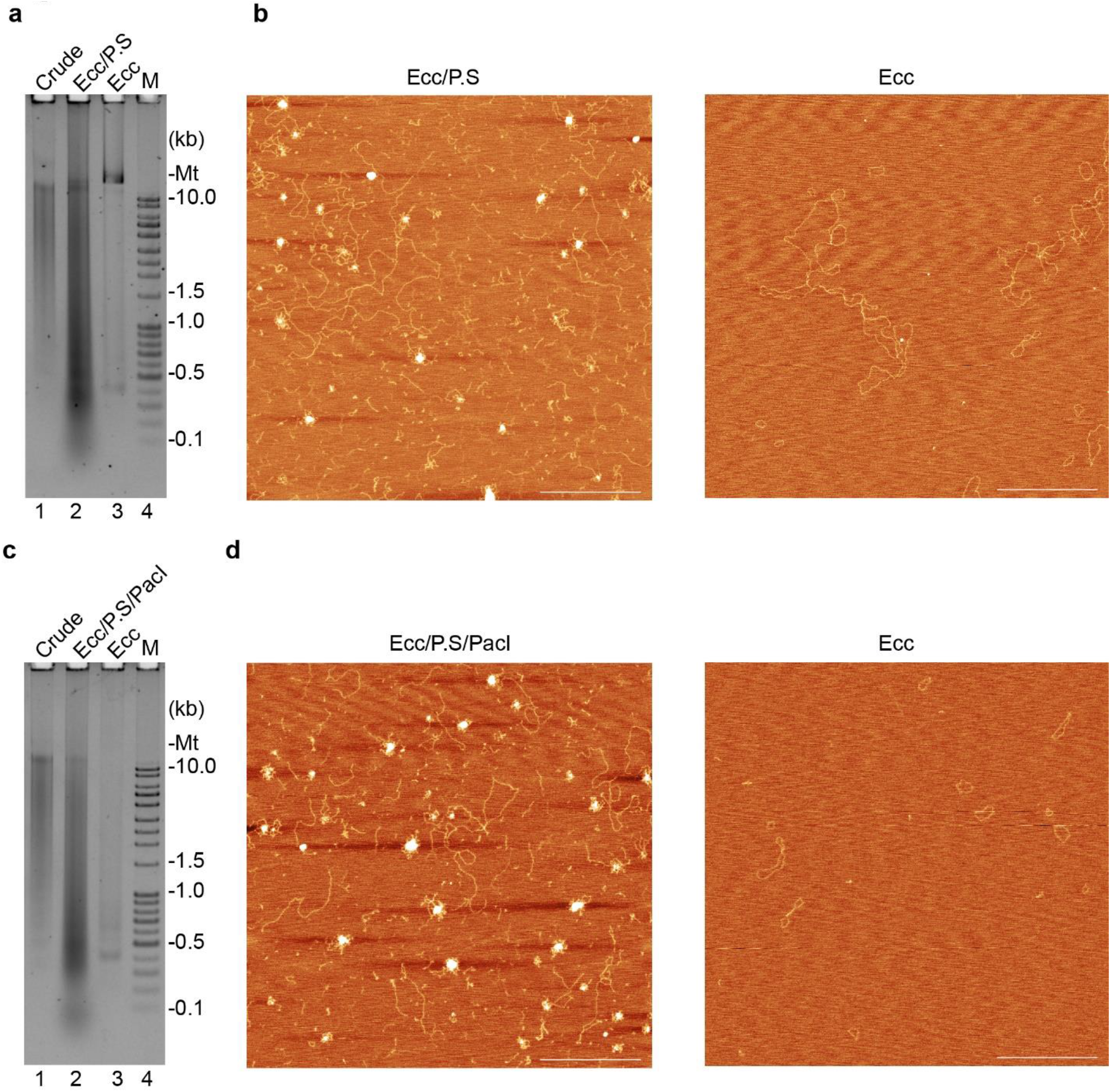
Agarose gel and SAFM images of eccDNAs at different purification steps. HeLa Cells were cultured in DMEM supplemented with 10% FBS, cells were continue cultured for another 48 hours after reaching 100% confluent; both floating and adherent cells were collected, washed with PBS for three times, resuspended in PBS and counted. Cells were fixed by adding absolute methanol to final 95% and put on ice for at least 10 minutes. Then 3SEP procedures were performed without PacI (**a-b**) or with PacI (**c-d**) linearizing mtDNA. **a**, Agarose gel image of the DNAs from different steps of 3SEP without PacI linearizing mtDNA. Lane 1: crude circular DNAs (Crude) after alkaline lysis from Step 1 #13; Lane 2: eccDNA after Plasmid-Safe DNase treatment (Ecc/P.S) from Step 2#18; Lane 3: eccDNA (Ecc) after Solution A purification from Step3 #33; Lane 4: linear DNA ladder; mtDNA (Mt); **Note**: To avoid over-loading, DNAs were not proportionally loaded. **b**, Atomic force microscopy image of the DNAs in Lane 2 (left panel, Ecc/P.S) and Lane 3 (right panel, eccDNA) in (**a**). The two big circular DNAs on the right panel are mtDNA purified by 3SEP in the absence of PacI. **c**, Agarose gel image of the DNAs from different steps of 3SEP with PacI linearizing mtDNA. All the information is the same as panel a except PacI is used in Step 2#18 (Lane 2). **d**, Atomic force microscopy image of the DNAs in Lane 2 (left panel, Ecc/P.S/PacI) and Lane 3 (right panel, eccDNA) in (**c**). Bar size =500 nm.

Researchers with basic training in biochemistry and molecular biology can easily implement this protocol. However, if the final eccDNA yield is less than 10 ng, we strongly suggest confirmation of the eccDNA purity by imaging using atomic force microscopy or electron microscopy. In this case, relevant trainings in sample preparation, microscopy, or assistance from relevant facilities are needed. To obtain the full-length sequence of eccDNA by Nanopore long read sequencing, basic computational skills or training provided by Oxford Nanopore are required for setting up the sequencing platform in a laboratory. In addition, basic Linux and Python skills are needed for processing the long reads and constructing full-length sequence of eccDNA by running the packages, which are written in Python.

### • Limitations

Although circular DNA with size bigger than 16 kb (mitochondrial DNA) could be efficiently purified using our method, the Nanopore sequencing method we described here can mainly process eccDNAs shorter than 10 kb. For eccDNAs longer than 10 kb, please refer to other long sequencing methods.

## REAGENTS LIST

- Methanol HPLC Grade (EMD Millipore MX0475-1)
- Pyrrolidine, 99% (Sigma-Aldrich P73803-100ML)
- Sodium Dodecyl Sulfate, 20% Solution (Fisher Chemical, BP1311200)
- 2-Mercaptoethanol ≥99.0% (Sigma-Aldrich, M7522)
- QIAGEN Plasmid Plus Midi Kit (25) (Qiagen, 12943)
- Plasmid-Safe™ ATP-Dependent DNase (Lucigen, E3110k)
- Solution A of cccDNA Purification Kit from either Bingene (220501-50) Or Biofargo (220501)
- Qubit™ 1X dsDNA HS Assay Kit (Thermo Scientific, Q33231)
- Phase Lock Gel™, QuantaBio, Phase Lock Gel Heavy (VWR, 10847-802)
- Phenol/Chloroform/Isoamyl Alcohol (25:24:1 Mixture, pH 8.0) (Thermo Fisher Scientific, BP1752I-400)
- Ethyl alcohol, Pure, 200 proof, for molecular biology (Sigma-Aldrich, E7023)
- 1 M Tris pH 7.0 (Fisher Scientific AM9851)
- UltraPure™ DNase/RNase-Free Distilled Water (Thermo Fisher Scientific,10977-023)
- SYBR™ Gold Nucleic Acid Gel Stain (10,000X Concentrate in DMSO) (Fisher Scientific, S11494)
- TE buffer (Thermo Fisher Scientific, AM9858)
- GLYCOGEN, MB GRADE 20 MG (Sigma-Aldrich, 10901393001)
- Dynabeads™ MyOne™ Silane (Thermo Fisher Scientific, 37002D)
- PacI (New England Biolabs, R0547)
- UltraPure UltraPure Agarose (Thermo Fisher Scientific, 16-500-100)
- EDTA (0.5 M), pH 8.0 (Thermo Fisher Scientific, AM9260G)
- Ultra-0.5 Centrifugal Filter Unit with Ultracel-10 membrane (Amicon, UFC5010)
- 5 M NaCl (Thermo Fisher Scientific, AM9760G)
- Sodium Acetate (3 M), pH 5.5 (Thermo Fisher Scientific, AM9740)
- Thermo Scientific Exo Resistant Random Primer (Thermo Fisher Scientific, SO181)
- phi29 DNA Polymerase (New England Biolabs, M0269)
- Deoxynucleotide Solution Set (New England Biolabs, N0446)
- Pyrophosphatase, Inorganic (New England Biolabs, M2403)
- BSA, Molecular Biology Grade (NEB, B9000S)
- SPRIselect (Beckman Coulter, B23317)
- T7 Endonuclease I (New England Biolabs, M0302)
- Ligation Sequencing Kit (Oxford Nanopore SQK-LSK109)

## EQUIPMENT LIST

- (Optional) DNA LoBind Tube 1.5 mL, PCR clean (Fisher Scientific, 13-698-791)
- QIAvac 24 Plus
- Thermal cycler
- Nanodrop (Thermo Scientific)
- Magnet (compatible with 1.5 ml tube)
- Qubit Fluorometer (Fisher scientific)
- Benchtop centrifuge
- Mini-PROTEAN(r) Tetra Vertical Electrophoresis Cell for Mini Precast Gels, 4-gel (Biorad, 1658004)
- Mini-PROTEAN(r) Tetra Cell Casting Module (15-well, 1.5 mm gel) (Biorad, 1658022)
- (Optional) Highest Grade V1 AFM Mica Discs (Fisher Scientific, NC1535937)
- (Optional) Dust-Off Disposable Air Duster (Falcon, DPSJB-12)
- Flow Cell (R9.4.1) (Oxford Nanopore, FLO-MIN106D)
- Sequencer (Oxford Nanopore)
- **Computational hardware**: a computer/server with at least 32GB memory and 8-core CPU running 64-bit Linux or macOS system. If the analysis starts from Nanopore raw data, a computer/server with NVIDIA GPU (NVIDIA compute version of 6.1 or higher) would significantly reduce the base-calling time.

## SOFTWARE LIST

- Guppy: base-calling from Nanopore raw signals. Registration is required to download. (https://community.nanoporetech.com/downloads)
- eccDNA calling toolkit (https://github.com/YiZhang-lab/eccDNA_RCA_nanopore)
- minimap2^20^ (https://github.com/lh3/minimap2)
- Porechop (https://github.com/rrwick/Porechop)
- Snakemake^21^ (https://snakemake.github.io)
- Git (https://git-scm.com/downloads)
- Python3 (https://www.python.org/downloads/)
- R (https://cran.r-project.org)
- Python packages:
  - pyfaidx (https://pypi.org/project/pyfaidx/)
  - pyfastx (https://pypi.org/project/pyfastx/)
  - Biopython (https://biopython.org)
- R packages:
  - caTools (https://cran.r-project.org/web/packages/caTools/)
  - data.table (https://cran.r-project.org/web/packages/data.table/)
  - digest (https://cran.r-project.org/web/packages/digest/)
  - dplyr (https://cran.r-project.org/web/packages/dplyr/)
  - DT (https://cran.r-project.org/web/packages/DT/)
  - emojifont (https://cran.r-project.org/web/packages/emojifont/)
  - extrafont (https://cran.r-project.org/web/packages/extrafont/)
  - fastmatch (https://cran.r-project.org/web/packages/fastmatch/)
  - flexdashboard (https://cran.r-project.org/web/packages/flexdashboard/)
  - futile.logger (https://cran.r-project.org/web/packages/futile.logger/)
  - ggplot2 (https://cran.r-project.org/web/packages/ggplot2/)
  - ggExtra (https://cran.r-project.org/web/packages/ggExtra/)
  - ggridges (https://cran.r-project.org/web/packages/ggridges/)
  - knitr (https://cran.r-project.org/web/packages/knitr/)
  - optparse (https://cran.r-project.org/web/packages/optparse/)
  - plyr (https://cran.r-project.org/web/packages/plyr/)
  - RColorBrewer (https://cran.r-project.org/web/packages/RColorBrewer/)
  - readr (https://cran.r-project.org/web/packages/readr/)
  - reshape2 (https://cran.r-project.org/web/packages/reshape2/)
  - scales (https://cran.r-project.org/web/packages/scales/)
  - tufte (https://cran.r-project.org/web/packages/tufte/)
  - viridis (https://cran.r-project.org/web/packages/viridis/)
  - yaml (https://cran.r-project.org/web/packages/yaml/)

## SOFTWARE SETUP

To install software listed above, the link for each software provides installation instructions. If the software is installed in a user specific path, this path should be added to the `$PATH` environment variable with shell command line export PATH=*/path/to/software/bin*:$PATH. To setup python packages, pip install *package_name* can be used. To install R packages, install.packages(*package_name*) can be used inside R.

Alternatively, software installation can be done by using the conda package management system. Instructions to setup conda are available at https://bioconda.github.io/user/install.html#install-conda. After setting up conda, software and packages can be installed with `conda install` `package_name`.

## EXAMPLE DATA

We provided an example dataset of raw sequencing signals in fast5 files generated by Nanopore sequencing of rolling-circle amplified eccDNA purified from mESC. These fast5 files are the raw input for the whole eccDNA calling pipeline. The dataset was subsampled from a whole run of MinION R9.4.1 Flow Cell (Oxford Nanopore, FLO-MIN106D) with sequencing kit (Oxford Nanopore, SQK-LSK109) in the original data in ref^16^. The example dataset can be downloaded at https://figshare.com/articles/dataset/Nanopore_reads_of_eccDNA/17046158.

### Reagent setup

#### Suspension buffer

10 mM EDTA pH8.0

150 mM NaCl

1% glycerol

Lysis blue (1000×, from Qiagen Kit) RNase A (110 mg/ml, 200×)

Keep at 4 °C. Freshly add 1/500 volume 2-Mercaptoethanol before use.

#### Pyr buffer

0.5M pyrrolidine 20 mM EDTA

1% SDS

Adjust pH to 11.80 with 2 M Sodium Acetate pH 4.00

Freshly add 1/500 volume 2-Mercaptoethanol before use.

#### (Optional) 10× AFM imaging buffer

100 mM NiCl_2_

100 mM Tris-HCl, pH 8.0

#### (Optional) 500× AFM sampling washing buffer

1M Magnesium acetate

#### 50X stock of TAE

Tris-base: 242 g

Acetate (100% acetic acid): 57.1 ml

EDTA: 100 ml 0.5 M sodium EDTA

Add dH2O up to one liter.

## Procedure

### EccDNA purification ●Timing 1-2 days

#### Step 1: Crude circular DNA isolation ●Timing 1.5 - 2 hours

To make this protocol easy to follow, here we present the eccDNA purification by 3SEP with over-confluent HeLa cells, which can be achieved by culturing cells in DMEM with 10% FBS for another 48-72 hours after cell reaching 100% confluent. We suggest to start with two 10 cm dishes of cells, and the following procedure is described for cells of about 35 million. Given eccDNA abundance is highly variable among biological samples and cellular status, the researchers need to determine the cell numbers required for eccDNA purification if different cell types and culture conditions are used.

1. (Optional) Fix cells. Collect both floating cells and adherent cells (by trypsinization). Wash cells with PBS for 3 times, count and resuspend cells with 0.5 ml PBS, then add 9.5 ml absolute methanol, keep on ice for 10 minutes. **PAUSE POINT** Fixed cells could also be stored at -20 °C for months.
2. Collect 35 million HeLa cells to a 50 ml tube, then centrifuge for 10 minutes with 2,000g at 4 °C.
3. Resuspend cells in suspension buffer. Suspend cells with 10 ml **Suspension buffer** after supplementing 20 μl 2-Mercaptoethanol. (for more cells, proportionally scale up) Note Ratio of suspension buffer volume to the cell number may vary according to cell properties. Low ratio of suspension buffer to cells will result in incomplete precipitation of SDS-potassium-DNA complex in Step 1 #7.
4. Add 10 ml **Pyr buffer (**equal volume of Step 1 #3) after supplementing 20 μl 2-Mercaptoethanol, gently mix by inverting tube for 5-10 times, lysate will become blue, keep 5 minutes at room temperature. **! CAUTION Pyr buffer** contains pyrrolidine, has an ammoniacal odor, should be handled in a fume hood or under a benchtop exhaust snorkel.
5. Add 10 ml **Buffer S3** (from QIAGEN Plasmid Plus Midi Kit, equal volume of Step 1 #3), gently invert tube until the solution color turns white.
6. Centrifuge the lysate for 10 minutes at 4,500 g. Note If the white SDS-potassium-DNA complex is not well precipitated, increase the ratio of suspension buffer volume to input cells.
7. Transfer the clear lysate to QIAfilter Cartridge (QIAGEN Plasmid Plus Midi Kit), and filter the cell lysate into a 50 ml tube.
8. Estimate the volume of filtrated lysate, add 1/3 of the estimated volume Buffer BB (QIAGEN Plasmid Plus Midi Kit). Mix by inverting tube for 4-8 times.
9. Transfer the lysate to the spin column on Qiavac 24 plus, apply vacuum until all liquid drawn through the column, according to Qiagen’s instructions.
10. Wash the column by 0.7 ml ETR buffer (QIAGEN Plasmid Plus Midi Kit), apply vacuum until all liquid drawn through the column. Repeat the wash with 0.7 ml PE buffer.
11. Centrifuge the column at 10,000 g for 2 minutes and transfer the column to a clean 1.5 ml tube.
12. Elute the crude circular DNA by adding 100-200 μl 0.1X EB buffer to the column, wait > 2 minutes and centrifuge at 10,000 g for 1 minute.
13. Measure the DNA concentration by either nanodrop or Qubit™ 1X dsDNA HS Assay Kit. **PAUSE POINT** DNA can be kept at 4°C overnight or -20°C for long-term storage.

#### Step 2: Digest linear DNA ●Timing 3.5 hours-overnight

14. Linearize mtDNA by PacI and digest linear DNA with ATP-dependent Plasmid-Safe DNase (Proportionally scale up the reaction according to the real amount of DNA from Step 13)

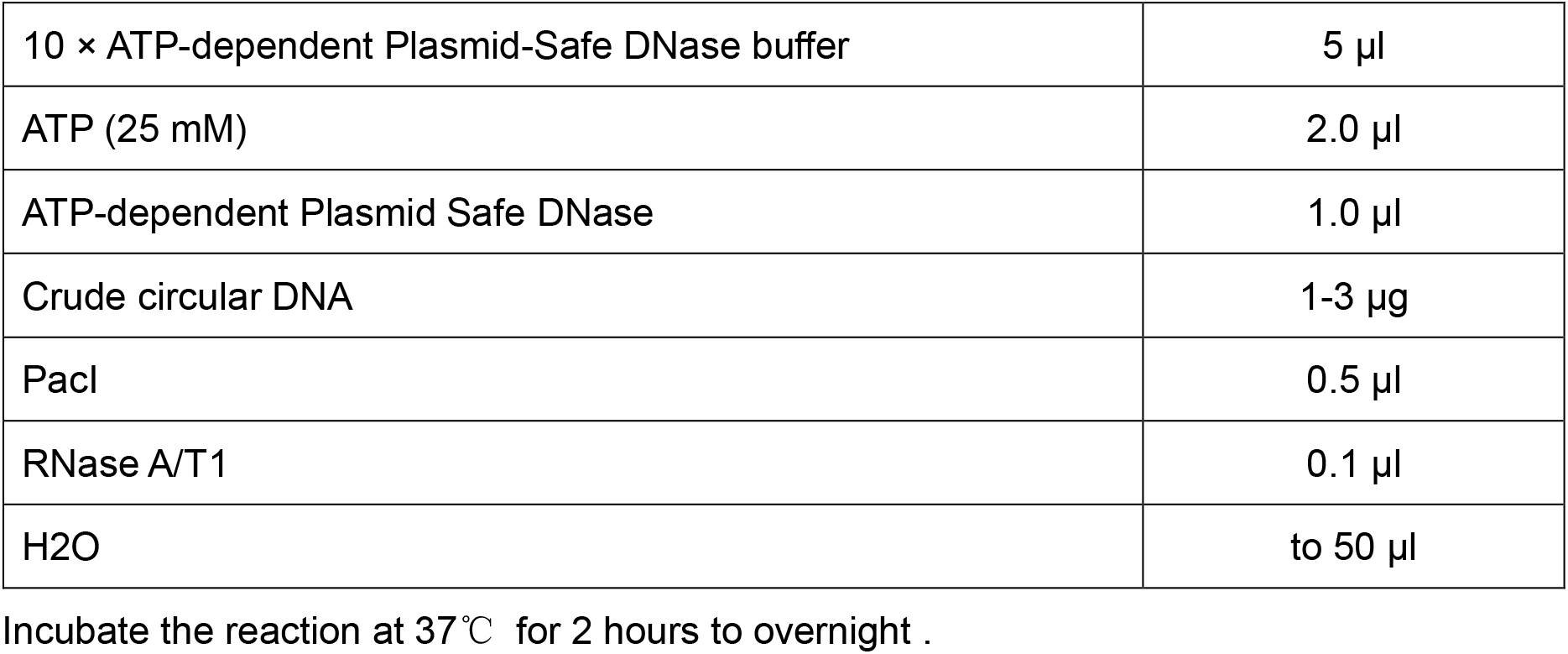 Note we use PacI to linearize mtDNA (both human and mouse mtDNA have 3 PacI recognition sites) here because it’s a rare cutter (8-bp recognition site) and is also fully active in the Plasmid-Safe DNase buffer. Thus, linearizing mtDNA and removing linear DNA could be performed in a single step. However, if PacI digestion is not desired, it could be omitted without any additional changes. Moreover, PacI could be replaced by any other rare cutters, or even CRISPR/Cas9 with sgRNA targeting mtDNA or any unwanted circular DNA. However, if the new rare cutters are not active in the Plasmid-Safe DNase buffer, the linearization must be performed before Plasmid-Safe DNase digestion, and proper buffer exchanging must be done before Plasmid-Safe DNase treatment.
15. (Optional) Concentrate solution. Filtrate the reaction solution from Step 14 by Ultra-0.5 Centrifugal Filter Unit (10 kDa) to ∼ 360 μl according to the manufacturer’s instruction.
16. Adjust the solution volume from Step 14 or Step 15 to 360 μl, then extract DNA with PCI. Transfer the solution from Step 14 or concentrate from Step 15 to Phase Lock Gel tube, add equal volume of Phenol/Chloroform/Isoamyl Alcohol (25:24:1 Mixture, pH 8.0), shake the tube by hand thoroughly and centrifuge at 14,000 g for 7.5 minutes. **! CAUTION** Phenol should be handled in a fume hood or under a benchtop exhaust snorkel.
17. Precipitate DNA. Transfer the aqueous phase to a new tube, add 1/10 volume of sodium acetate (3 M, pH 5.5), add 1 μl glycogen and 3 volume of 200 proof ethanol, mix and put at - 80 for at least 30 minutes. **PAUSE POINT** DNA could be kept at -80°C for overnight or longer.
18. Centrifuge the DNA for 30 minutes at 4 °C, wash the pellet once with 1 ml 80% ethanol. Spin down the tube, pipet out residual ethanol, resuspend the pellet with 50 μl 2 mM Tris-HCl 7.0.

#### Step 3: Selectively recover eccDNA ●Timing 1-1.5 hours

19. Equilibrate Solution A at room temperature for 30 min, add 700 μl Solution A and mix the solution by pipetting up and down for 10 times, and let it stand for 5 minutes at room temperature.
20. Resuspend the Dynabeads™ MyOne™ Silane beads by thoroughly vortex, take 10 μl to a new tube and still it on appropriate magnet until all bead settling aside, remove the liquid and resuspend the beads in 20 μl Solution A. **! CAUTION** Solution A contains phenol and should be handled in a fume hood or under a benchtop exhaust snorkel.
21. Combine the suspended beads with solution from Step19 and pipetting up and down for 10 times, incubate at RT for 5 minutes.
22. Place the tube on a magnet and allow the beads to settle, remove and discard the solution (circular DNAs are on beads now).
23. Quickly spin down the tube using benchtop centrifuge for 10 second and put it on magnet again, remove the residual liquid with 10 μl pipet tips. ▲ **CRITICAL STEP** The less free-liquid left, the purer eccDNA will be, but with caution not to remove the beads, which are bound with circular DNA.
24. Take off the tube from magnet, resuspend the beads in 300 μl Solution A by pipetting, stand the tube for 2 minutes.
25. Place the tube on magnet and allow the beads settle to magnet, remove and discard the solution.
26. Quickly spin down the tube using benchtop centrifuge for 10 second and put it on magnet again, remove the residual liquid with 10 μl pipet tips. ▲ **CRITICAL STEP** The less free liquid left, the purer eccDNA will be, but with caution not to remove the beads, which are bound with circular DNA.
27. Repeat Step 24-26 one more time.
28. Keep tube on magnet, add 700 μl 3.5 M NaCl and wait for 1 minute without disturbing the beads, remove the solution. Repeat once.
29. Keep the tube on magnet, add 800 μl freshly prepared 80% ethanol, wait for 1 minute without disturbing the beads, remove the ethanol. Repeat once.
30. Quickly spin down the tube using benchtop centrifuge for 30 seconds, and put the tube on magnet, remove all residual liquid with 10 μl pipet tips with caution not to remove beads.
31. Take off the tube from magnet, thoroughly resuspend the beads with 100 μl 0.1 X EB buffer before beads are completely dried out, rotate the tube slowly on a rotator for > 3 minutes.
32. Place the tube on magnet to settle down the beads. Transfer the elute to a new DNA LoBind Tube.
33. Measure the eccDNA concentration. Given the low abundance of eccDNA, we usually use Qubit™ 1X dsDNA HS Assay kit to measure the eccDNA concentration. In general, at least 10 ng pure eccDNA could be obtained from 35 million over-confluent HeLa cells. Note: Note: Given eccDNAs content in most biological samples was low and their precise mass quantification always requires highly sensitive methods. However, we found commercially available fluorescence dye-based sensitive kits, including Qubit™ 1X dsDNA HS Assay kit, underestimates the concentration of eccDNAs of hundreds bp by 1-4 folds in a circle-size dependent manner. To precisely quantify eccDNAs of low concentration, we developed a SYBR gold-based quantification method that could precisely quantify both pure linear dsDNA and circular dsDNA with identical standard curve generated by linear DNA. Briefly, quantification working solution is prepared by diluting SYBR gold dye concentrate with 1x TE buffer (10 mM Tris 8.0, 1 mM EDTA) at ratio of 1:10000; DNA from Qubit™ 1X dsDNA HS Assay kit is used to generate the standard curve for quantification of both linear DNA and eccDNAs; then DNA are mixed with quantification working solution to 100 ul and fluorescence densities are read in a black 384 wells plate with plate reader at Ex/Em= 495-8/536-11 nm, DNA concentrations are calculated according to the standard curve after linear regression.

#### Vertical agarose gel electrophoresis. ●Timing 2 hours

34. Mix 1.0 g ultra-pure agarose with 100 ml 1X TAE in a microwave flask, microwave for 3-5 minutes until the agarose is completely dissolved. Note Gel concentration may be varied from 1-2 %.
35. Pour the agarose to the pre-assembled glass plates immediately after the agarose is dissolved without cooling down. Insert the comb and let the gel cool down at least 20 minutes. **! CAUTION HOT!** Wear heat resistant gloves to handle the flask. Be careful during stirring as eruptive boiling can occur.
36. Remove the comb after dismounting gel from casting chamber, (optional) remove residual gel slices within wells by a sharp tweezer.
37. Assemble the gel to the apparatus according to the manufacturer’s instructions, then fill the chamber with 1X TAE.
38. Carefully pipet samples (> 1 ng eccDNA could be visualized) into wells, and load suitable amount of DNA ladder.
39. Separate DNA at 80V X 35 minutes, running time may be different depending on the gel concentration in Step 34.
40. Turn OFF power, open the glass plates and transfer the gel to a 15 cm diameter plastic dish, add 50-70 ml 1X TAE.
41. Add 5 μl SYBr Gold concentrate, shake gel for > 15 minute in dark.
42. Visualize the gel using a gel imaging device that has blue light.

#### (Optional but recommended) Scan atomic force microscopy imaging. ●Timing ∼2 hours

43. Take 4.5 μl eccDNA with a concentration at 0.6-1.0 ng/μl, add 0.5 μl (1/10 volume) 10× imaging buffer and mix by pipetting.
44. Cleave mica surface by double side tape, spread DNA mixture on the mica surface and incubate for 2 min.
45. Blow off the liquid with compressed gas, rinse the mica twice with 30 μl of 2 mM magnesium acetate. Repeat once.
46. Acquire image by using tip C of an SNL-10 probe and scanning in Air mode and processing with Gwyddion. (e.g. scanned with Veeco MultiMode atomic-force microscope with a Nanoscope V Controller in ‘ScanAsyst in Air mode’)

#### Rolling cycle amplification and Nanopore sequencing library construction ●Timing 2 days

47. <u>Rolling cycle amplification (RCA) of eccDNA</u>

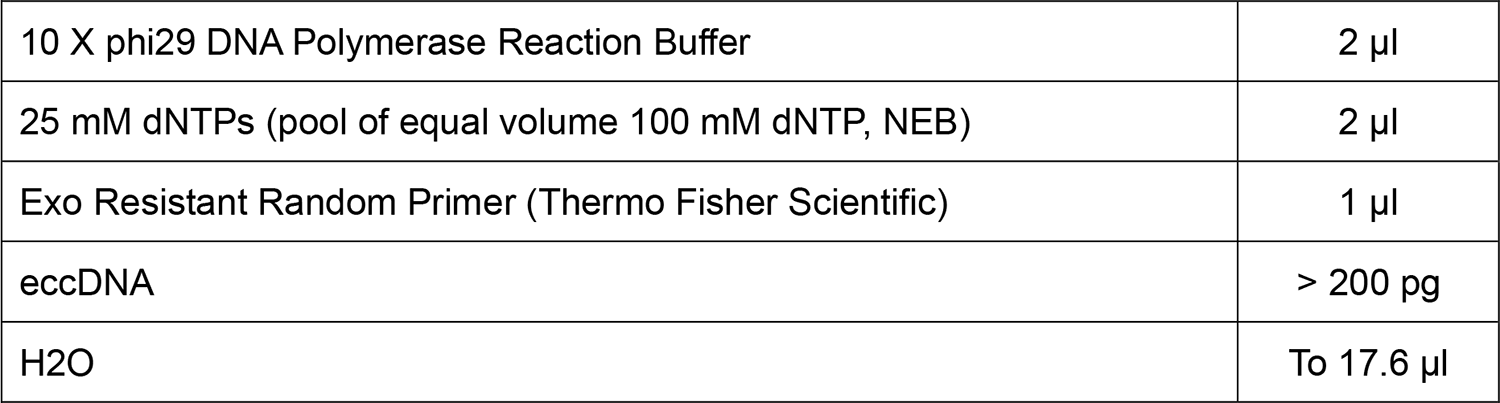 Incubate at 95°C for 5 minutes, then ramp to 30 °C at 1% ramp rate on thermocycler (∼30 minutes).

48. Dilute the RCA product with 180 μl ultra-pure water, recover the high molecular weight (HMW) DNA with 80 μl SPRIselect beads (0.4 X Volume) according to the manufacture’s instruction.
49. Debranch RCA products

Incubate at 95°C for 5 minutes, then ramp to 30 °C at 1% ramp rate on thermocycler (~30 minutes).
|  |  |
| --- | --- |
| phi29 DNA Polymerase (NEB) | 1 μl |
| Pyrophosphatase, Inorganic (NEB) | 1 μl |
| BSA, Molecular Biology Grade | 0.4 µl |

Incubate at 30°C for 14 hours, inactivate at 65°C for 10 minutes.

50. Add 210 μl elution buffer (Qiagen), immediately extract the debranched products with 360 μl Phenol/Chloroform/Isoamyl Alcohol (25:24:1 Mixture, pH 8.0). Then precipitate DNA as in Step 17. **! CAUTION** Phenol should be handled in a fume hood or under a benchtop exhaust snorkel.
51. Resuspend DNA with 200 μl as performed in Step 18.
52. Select the high molecular weight DNA with 80 μl (0.4X V) SPRIselect beads as in Step 48.
53. Construct Nanopore sequencing library with Ligation Sequencing Kit (Oxford Nanopore) according to the manufacture’s instruction.

|  |  |
| --- | --- |
| 10X NEBuffer 2 | 15 µl |
| T7 Endonuclease I | 7.5 µl |
| (HMW) DNA | 3 µg |
| H2O | To 150 µl |

Incubate at 37°C for 15 minutes.

#### Nanopore sequencing ●Timing 2 days

54. Perform the Nanopore long-read sequencing with appropriate flow cell (e.g. FLO-MIN106 for R9.4.1 flow cell) according to the manufacture’s instruction. ●Timing 2 days

#### Data processing and analysis ●Timing 1 day to 1 week

1. Download the reference genome sequences of the studied species in fasta format from NCBI, EMBL-EBI, UCSC genome browser or Ensembl. For example, mouse reference genome (GRCm38 / mm10) can be obtained by:

~~~
wget -O GRCm38.fa.gz
https://ftp.ebi.ac.uk/pub/databases/gencode/Gencode_mouse/release_M25/GRCm38.primary_assembly.genome.fa.gz
gzip -d GRCm38.fa.gz
~~~
2. Build minimap2 index of the reference genome.

~~~
minimap2 -x map-ont -d GRCm38.mmi GRCm38.fa
~~~
3. Create a working folder and download eccDNA analysis scripts.

~~~
mkdir eccDNA
cd eccDNA
git clone https://github.com/YiZhang-lab/eccDNA_RCA_nanopore.git
~~~ To run the example data, use the following commands to download the data:

~~~
wget –O nanopore_fast5.zip https://figshare.com/ndownloader/files/31526759
unzip nanopore_fast5.zip
rm nanopore_fast5.zip
~~~

##### Base calling and mapping

4. The sequencing data processing and mapping can be done using a single Snakemake based script with configuration file (option A) or run step-by-step commands (option B).

The output of an Oxford Nanopore sequencing run is the raw signals stored in multiple fast5 files. `guppy` will be used to perform base calling, which would convert the raw signals to DNA sequences stored in fastq format.

!CAUTION `guppy` base calling supports running in either CPU or GPU. However, running in CPU would be very time-consuming. Using a CUDA enabled GPU would greatly reduce the base calling time.

##### (A) Base calling and mapping using a single script

(i) The `eccDNA_RCA_nanopore/mapping/Snakefile` is the snakemake based script to perform base calling and reads mapping. Before running it, the configuration file `config.json` should be properly configured, with any text editor.

~~~
cd eccDNA_RCA_nanopore/mapping
vi config.json
~~~ Edit the parameters in the `config.json` according to the guidance in Table 1.

**Table 1.** Description of parameters in the configuration file.
(ii) Start the script with the following command. The `-j` option specifies the number of threads used in this run, which should be adjusted based on available CPU/GPU resources.

| Parameter | Description |
| --- | --- |
| sample | Sample name |
| fast5 | The absolute path to the folder of fast5 files generated by the Nanopore sequencer |
| flowcell | The Nanopore flow cell used when sequencing (e.g. FLO-MIN106 for R9.4.1 flow cell) |
| kit | The library construction kit used for sequencing (e.g. SQK-LSK109) |
| threads | Number of threads used in the run |
| reference | Path to the minimpa2 indexed reference genome ending with .mmi |
| workdir | Absolute path to the working folder |
| GPU | Whether GPU is available for base calling (Y: Yes, N: No) |

~~~
snakemake -j 8 –p
~~~ When the script finished running, the base calling results would be stored in the *sample_name*.fastq.gz file, and the mapping results are in the *sample_name*.paf file. The QC reports for the Nanopore sequencing run are provided under the *sample_name*_qc folder.

##### (B) Step-by-step base calling and mapping

(i) Use `guppy` to generated reads sequences from raw signals:

~~~
guppy_basecaller --input_path *<path/to/fast5/>* --save_path
*<path/to/output/folder>* --flowcell *flow_cell_name>* --kit
*<library_kit_name>* --calib_detect --num_callers *<threads>* --
trim_barcodes --trim_strategy dna --disable_pings --device auto
~~~ ▲CRITICAL The above command is for running with GPU. If GPU is not available, remove the `--device auto` option to run with CPU.
(ii) Trim adaptors from reads using `porechop`:

~~~
porechop --extra_end_trim 0 --discard_middle -i
*<guppy/output/pass/>* | gzip > *sample_name*.fastq.gz
~~~
(iii) Reads mapping with `minimap2`:

~~~
minimap2 -x map-ont -c --secondary=no -t *<threads>
path/to/reference.<mmi> sample_name*.fastq.gz > *sample_name*.paf
~~~ !CAUTION When using fastq files as input, skip step (i) and starts from step (ii).

##### eccDNA calling

5. Consensus eccDNA calling and full-length sequence reconstruction are performed with `eccDNA_RCA_nanopore.py` using *sample_name*.fastq.gz and *sample_name*.paf generated in the last step as input:

~~~
./eccDNA_RCA_nanopore.py --fastq mapping/*sample_name*.fastq.gz --paf
mapping/*sample_name*.paf --info *info.<tsv>* --seq *seq.<fa>* --var
*var.<tsv>* --reference *path/to/reference.<fa>* --verbose | tee *<out.log>*
~~~ Additional parameters that can be adjusted could be found by command: `./eccDNA_RCA_nanopore.py -h`.

##### Anticipated results

By omitting or including PacI in Step 2 of 3SEP, we purified mtDNA retained (**Fig. 2a**-**b**) and depleted (**Fig. 2c**-**d**) eccDNAs from over-confluent HeLa cells. Agarose gel electrophoresis revealed the stepwise DNA pattern changes when eccDNAs were enriched by 3SEP (**Fig. 2a** and **c**). In general, more than 95% of contaminating DNA present in the alkaline lysis produced crude extrachromosomal circular DNA (Crudes in Lane 1 of **Fig. 2a** and **c**) can be degraded by overnight Plasmid-Safe DNase treatment. However, the rest 5% contaminating linear DNAs that escaped the Plasmid-Safe DNase treatment (Lane 2 of **Fig. 2a** and **c**) are still dominate the eccDNA when viewed by microscopy imaging (left panels of **Fig. 2b** and **d**). Nevertheless, upon further purification by Step 3 of 3SEP, eccDNAs of high purity were obtained and linear DNAs are barely detected (right panels of **Fig. 2b** and **d)**. The results support the efficacy and robustness of 3SEP in eccDNA enrichment.

Then the purified eccDNAs will be subjected to rolling cycle amplification and sequencing library construction before sequencing. Then the sequencing data will be subjected to processing. After the eccDNA calling scripts are finished, four output files will be generated. The reconstructed eccDNA information would be stored in the output info.tsv file. Table 2 explains the format of the eccDNA calling results. Example outputs of consensus eccDNA calling are provided in Table 3. eccDNAs with full-pass number of at least 2 are high-confident eccDNAs and can be used for further analysis. It is possible that reads with single-pass could also derive from eccDNAs. However, they cannot be discriminated from amplifications of linear DNAs. Thus, to avoid false positives we suggest using a minimum of two full-passes to make sure that the eccDNAs called are real circular DNA. The full-length sequence of each eccDNA is available in the `seq.fa` output file in the fasta format. `var.tsv` contains the identified variants information, with 6 columns representing chromosome, position, reference nucleotides, alternative nucleotides, supportive coverage and total coverage, respectively. `out.log` stores the detailed reconstruction and identification process for each eccDNA (examples provided in Box 1).

**Table 2.**
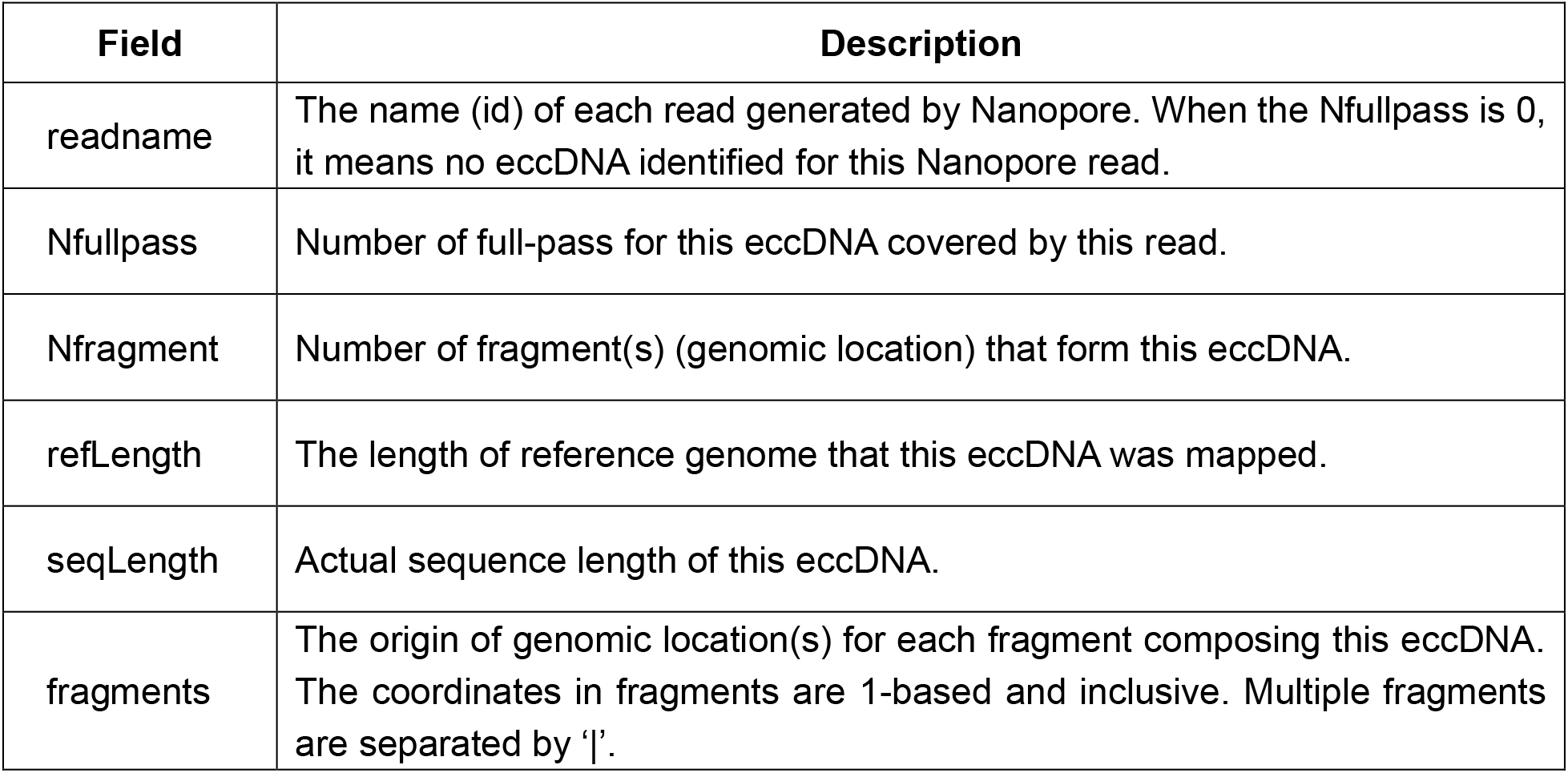
Description of eccDNA calling results in the info.tsv file.

**Table 3.**
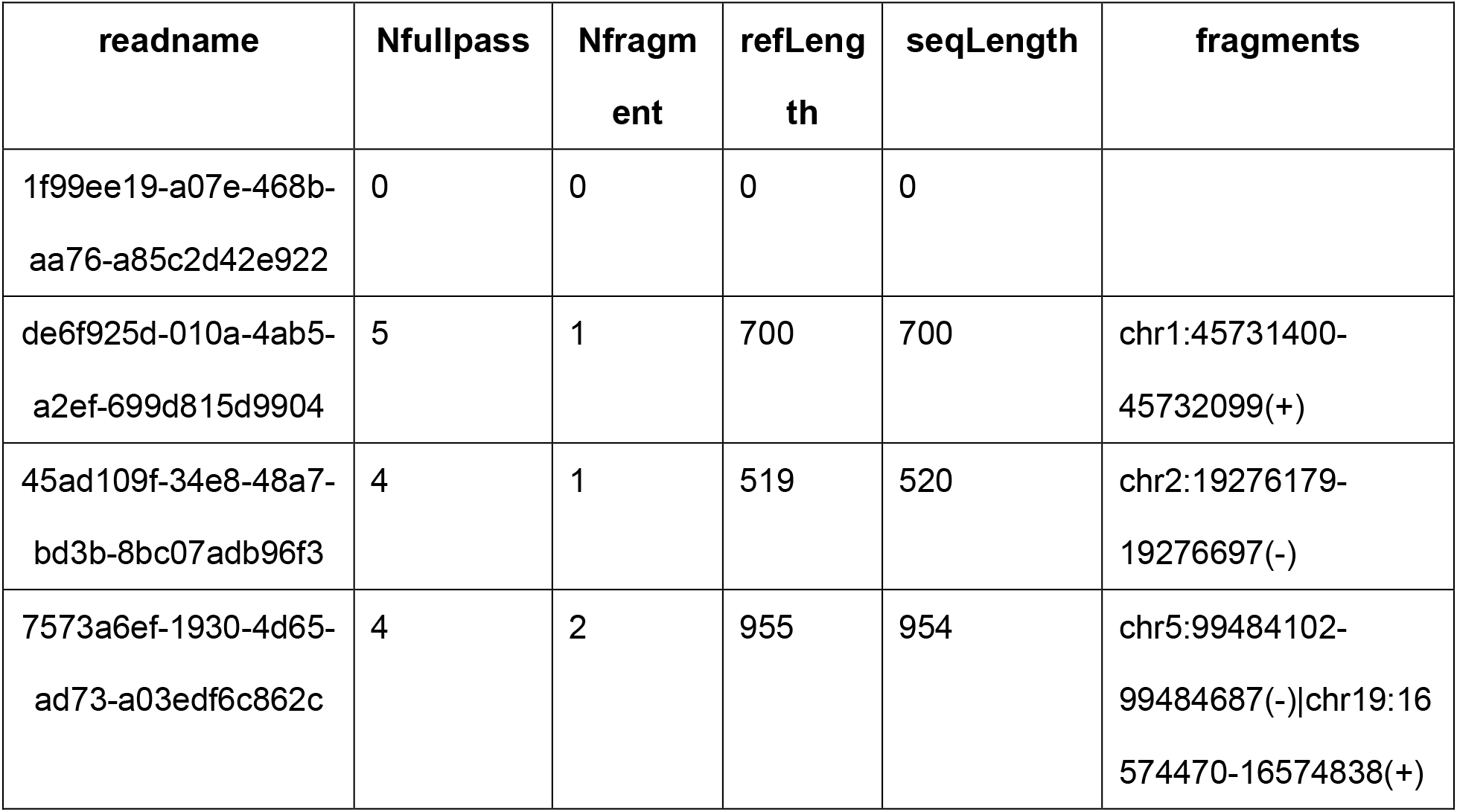
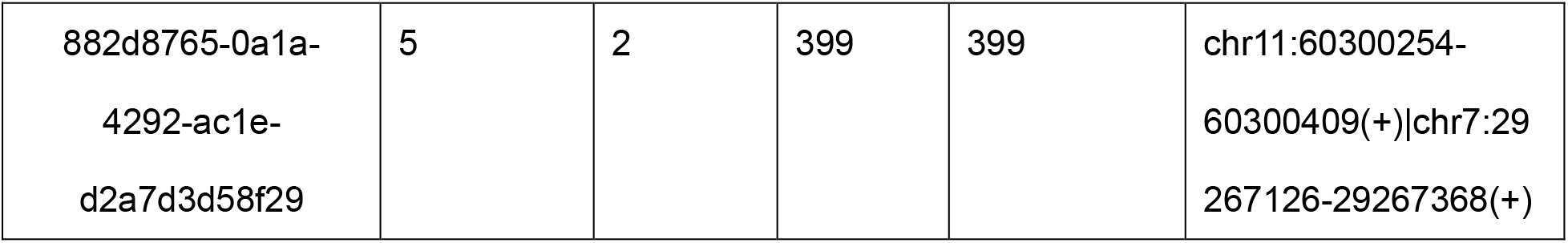
Example outputs of eccDNA calling results.

###### Box 1|

Example of detailed eccDNA reconstruction process from Nanopore reads of rolling-circle amplified eccDNA

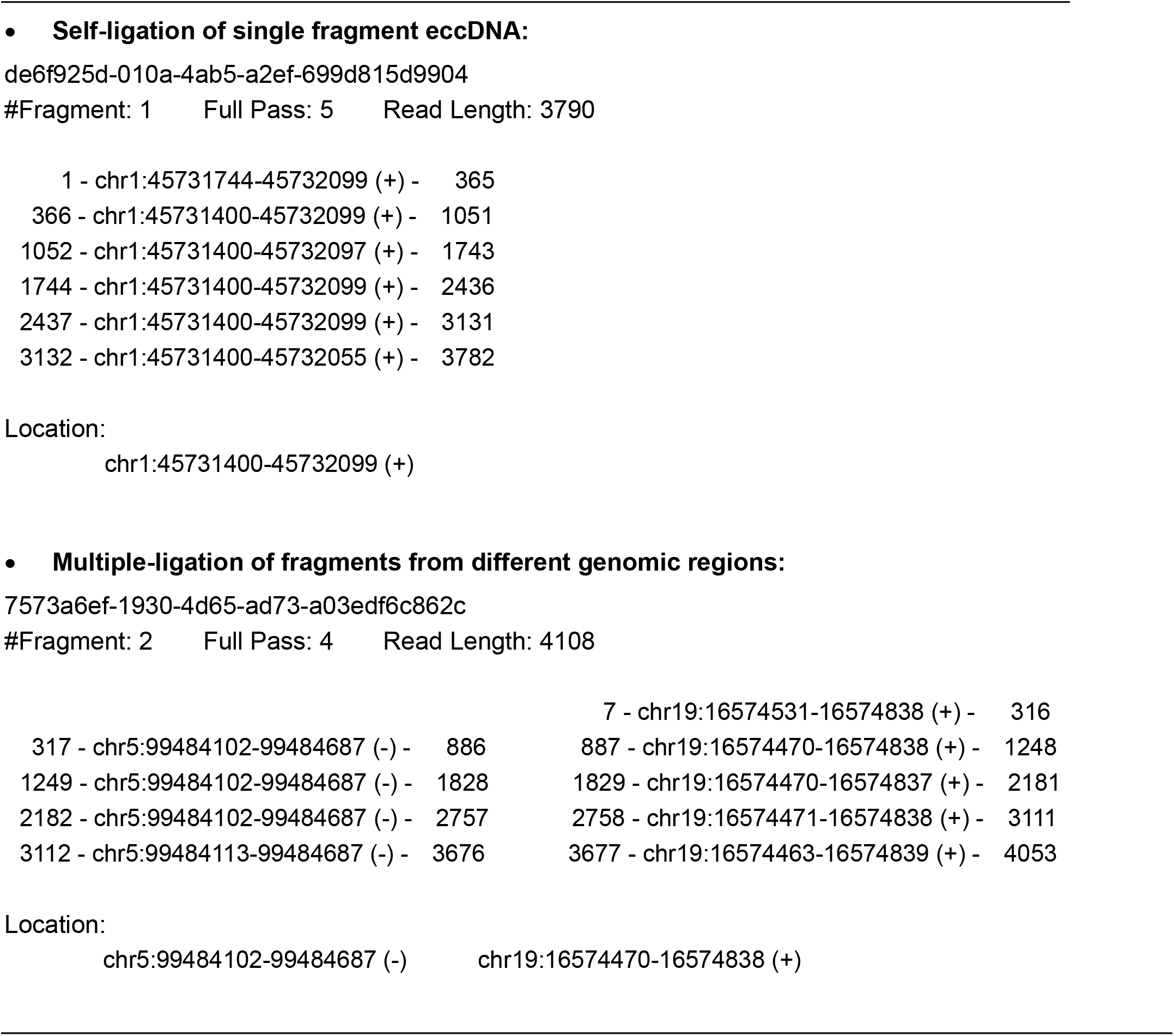

## Data availability

The example dataset of Nanopore sequencing of rolling-circle amplified eccDNA can be downloaded at https://figshare.com/articles/dataset/Nanopore_reads_of_eccDNA/17046158.

## Code availability

All scripts used for data analysis described in this paper are available at https://github.com/YiZhang-lab/eccDNA_RCA_nanopore.

## Notes

https://figshare.com/articles/dataset/Nanopore_reads_of_eccDNA/17046158

### Competing Interest Statement

The authors have declared no competing interest.

